# Differential Tissue Coupled Powering for Battery-Free Injectable Electroceuticals

**DOI:** 10.1101/2025.11.09.687502

**Authors:** Han Wu, Sultan Mahmud, Mali Halac, Domenica Baez, Jeremiasz Dados, Chaojun Cheng, Asif Iftekhar Omi, Anyu Jiang, Cameron Wallace, Anvi Singh, Erin Patrick, Baibhab Chatterjee, Shriya Srinivasan, Adam Khalifa

## Abstract

Electroceutical implants that deliver targeted neural stimulation have shown therapeutic potential for a wide range of neurological and peripheral disorders, yet wirelessly powering ultra-miniaturized, fully injectable systems remains a critical challenge. Here we report a Thread-like Injectable Neural TechnologY (TINY), powered via a differential tissue-coupled powering (DTCP) scheme. DTCP employs mid-frequency differential potentials applied across external electrodes on a compact, wearable transmitter to deliver energy through tissue to an ultra-miniaturized, thread-like implant that integrates a custom ASIC and PEDOT-coated receiver and stimulation electrodes. Benchtop experiments in agar phantoms characterize the power-transfer efficiency (PTE) and reveal that PTE increases with implant length while maintaining strong tolerance to angular misalignment. In vivo tests in rat hindlimbs further demonstrate wireless activation of the sciatic nerve through tissue at centimeter-scale depths, confirming effective transcutaneous energy delivery for neurostimulation. A 20-day implantation study shows stable positioning of the device with minimal tissue response, indicating excellent chronic compatibility. These findings address long-standing challenges in wirelessly powering injectable electroceuticals and establish DTCP as a scalable and alignment-robust powering strategy for future minimally invasive neuromodulation therapies.

## Introduction

Electroceuticals are gaining traction as therapeutic tools for disorders of the central and peripheral nervous systems. Within the central nervous system (CNS), spinal cord stimulation has demonstrated efficacy in treating chronic pain and movement impairments. In the peripheral nervous system (PNS), vagus, sacral, and tibial nerve stimulation have been used to manage inflammatory, gastrointestinal, and urinary disorders, while somatic nerve interfaces restore motor and sensory function in paralyzed or injured limbs 0. Collectively, these neural electroceuticals offer precise, reversible, and localized control of physiological functions, providing therapeutic alternatives where conventional pharmaceuticals are ineffective or poorly tolerated [2].

Current spine or peripheral nerve stimulators require open surgical implantation to accurately position leads on millimeter-scale nerves, ensuring stable contact and avoiding off-target activation [3]–[4]. Furthermore, these systems are bulky, necessitate invasive procedures and implanted pulse generators (IPGs), and are poorly suited for distributed or multi-site interfacing. The invasiveness and duration of these surgical procedures often relegate implantable electroceuticals to use only as a *last-line intervention* when conventional therapies fail.

As summarized in Fig. 1a, existing nerve stimulation strategies can be broadly categorized by their invasiveness, efficiency, and long-term reliability. (A) Transcutaneous electric nerve stimulation (TENS) provides excellent safety and noninvasiveness, but suffers from poor spatial precision and low coupling efficiency through tissue [5]. (B) Injectable wireless stimulators combine wireless powering with minimally invasive delivery, allowing small implants to be placed close to target nerves through a fine needle [6]–[11]. (C) Implantable wirelessly powered stimulators eliminate leads but still require pocket formation and precise surgical placement [12]–[19]. (D–E) Lead and cuff electrode systems integrate onboard batteries and wired electrodes, delivering the highest efficiency and chronic stability, but at the cost of extensive tunneling and high surgical invasiveness [20]–[22]. Surgical tunneling of electrodes and subcutaneous pulse generators introduces heavy clinical burden, while tethered connections restrict the placement and number of stimulation sites. As a result, the past decade has witnessed a shift toward wireless power transfer (WPT) technologies for implantable medical devices (IMDs). These approaches enable smaller form factors, reduced device weight, extended operational longevity, and, critically, less invasive implantation, thereby positioning categories (B-C) as the natural evolutionary directions for future systems.

**Fig. 1.**
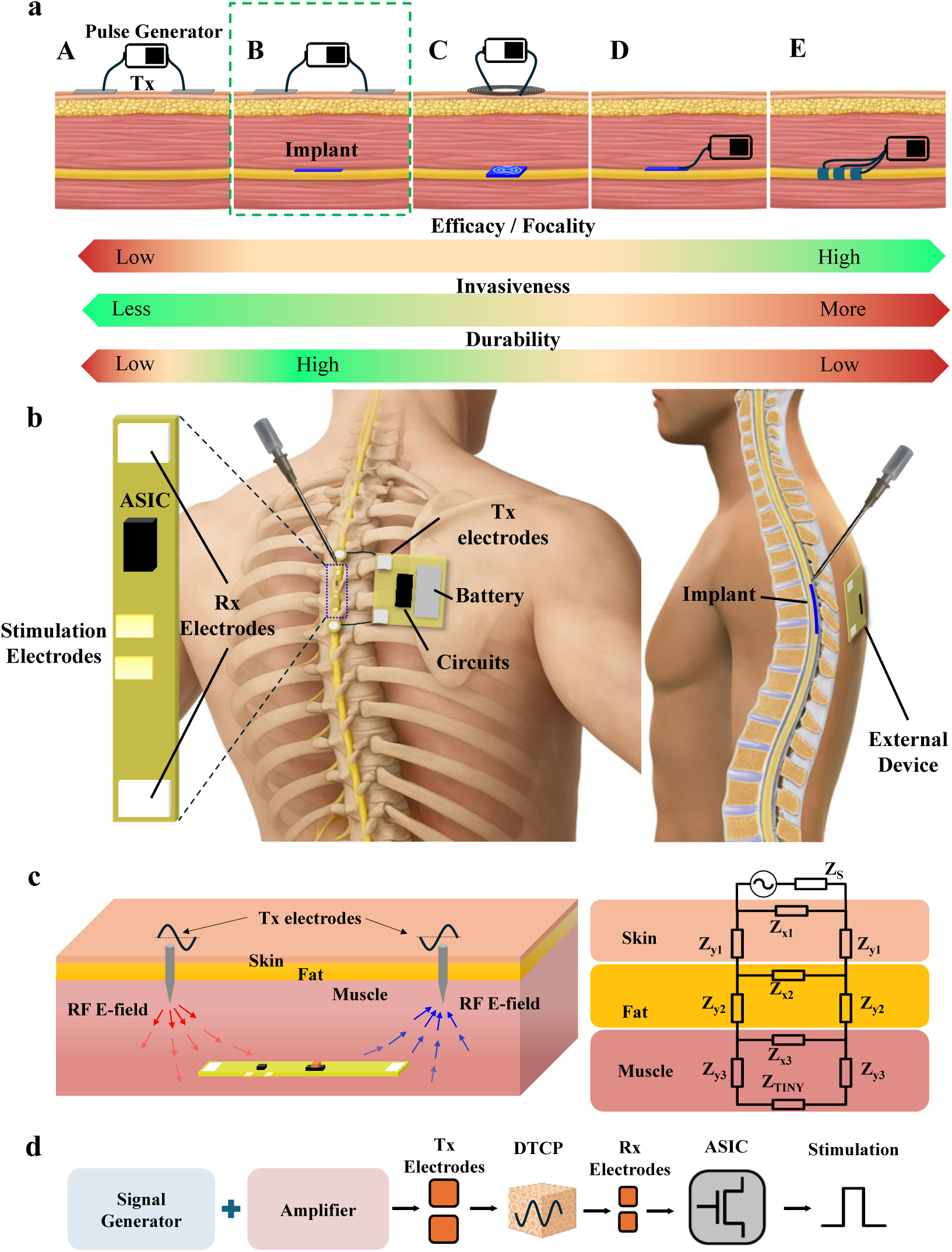
Overview of the DTCP injectable neural stimulator. **a.** Nerve stimulation powering methods comparison on efficacy/focality, invasiveness and durability. A. Transcutaneous electric nerve stimulation (TENS); B. Injectable wireless stimulator; C. Implantable wireless stimulator; D. Implanted lead stimulator; E. Implanted cuff electrodes. **b.** Concept of a TINY device injected onto the spinal nerve and powered by a portable transmitter device through DTCP. TINY device integrates receiver electrodes, stimulation electrodes and ASIC. **c.** Working principle of DTCP and a simplified tissue coupling circuit model. The implant’s receiver electrodes harvest the electric field from the transmitter, and the impedance of different tissue layers will impact the total efficiency. **d**. Block diagram of the overall electronic system, consisting of the external RF generator, tissue coupling path and the circuits on the implant to convert RF energy to DC stimulation.

However, while WPT represents a major step toward reducing surgical burden, achieving true minimal invasiveness requires not only eliminating leads and batteries but also eliminating open surgical dissection of pocket creation. The emerging concept of the injectable neurostimulator aims to meet this goal by delivering ultra-miniaturized stimulators directly into target tissues through a fine needle, avoiding open surgery and subcutaneous tunneling altogether. (B) Injectable systems, combining minimally invasive delivery with efficient and localized stimulation, offer the most balanced trade-off between safety, adaptability, and performance, making them the most promising direction for next-generation electroceutical systems.

Unfortunately, conventional WPT methods, such as inductive, ultrasonic, magnetoelectric, and optical coupling, face significant challenges in human-centric applications. Magnetoelectric and optical approaches, while offering good efficiency or precision, are inherently incompatible with injectable form factors. ME systems rely on resonant Metglas–PZT laminates whose required thickness and encapsulation substantially increase device volume [23][24], whereas optical links suffer from severe tissue scattering, limiting penetration to sub-millimeter depths with insufficient power delivery [25]–[28]. Ultrasonic links can wirelessly power millimeter-scale stimulators at centimeter depths benefiting from low acoustic attenuation in soft tissue [6]–[8]. However, the need for precise transducer alignment and coupling gel severely limits practical usability. The external acoustic heads remain bulky, and lateral misalignment quickly degrades coupling efficiency. As a result, it remains unsuitable for distributed or fully wearable injectable systems. Several groups have explored mid-field or inductively coupled RF powering to enable miniature, battery-free neurostimulators that approach the injectable form factor. *Neuspera* developed a mid-field inductively powered sacral nerve stimulator capable of activating deep neural targets in human trials [9][10]. However, the device remains too large (50 mm^3^). More recently, *Fuglevand et al.* presented a smaller ferrite-core coil stimulator (7 mm^3^) intended for injectable use [11]. While this design reduces the overall size, it remains rigid and mechanically incompatible with flexible implantation, and it is highly sensitive to angular alignment. Both studies also share a critical limitation, the absence of any quantitative characterization of power-transfer efficiency, highlighting the intrinsic challenges of achieving both efficient RF coupling and true injectability. Alternatively, *Injectrode* (Neuronoff Inc.) [29][30] employs a syringe-delivered conductive path to couple with transcutaneous stimulation, but its millimeter-scale wire and lack of integrated circuitry hinder scalability and future closed-loop functionality. While recent advances in wireless powering and miniaturized fabrication have enabled sub-millimeter and needle-deliverable implants in small animals, each existing modality remains constrained by inherent trade-offs between efficiency, depth, and form factor. As a result, current approaches still compromise adaptability, wearability, and overall user comfort, underscoring the need for a fundamentally different powering mechanism.

Our proposed strategy, Differential Tissue-Coupled Powering (DTCP), employs differential signal coupling through the body to deliver mid-frequency currents (~1–70 MHz) safely within established human exposure limits [31]. Non-invasive epidermal electrodes are positioned near the implant site to maximize power transfer via cavity resonance. A key innovation of this method lies in the *electronic rectification* of these benign alternating currents, which are transmitted through biological tissue via differential coupling [32]. The implant captures this energy and converts it into stable DC power to drive integrated circuits for stimulation. This mechanism enables the realization of sleek, battery-free devices suitable for minimally invasive injections. A few prototypes have demonstrated neural recording or stimulation via tissue volume conduction, using surface or textile electrodes coupled to implanted PCBs for EMG or EEG interfaces [33]–[37]. Despite proof of concept, these systems remain limited by bulky implant size, limited power transfer efficiency (PTE), and potential exceedance of IEEE C.95.1 [38] current exposure limits.

To translate the DTCP concept into a practical electroceutical platform, we develop the first fully injectable neural stimulator that implements DTCP end-to-end—from external transmission to subcutaneous energy capture and stimulation, termed Thread-like Injectable Neural TechnologY (TINY) as shown in Fig. 1b. TINY overcomes the limitations of previous designs through three key innovations: First, the entire system, including the application-specific integrated circuit (ASIC), power-receiving electrodes, and stimulation electrodes, is built into a single flexible, strand-like form factor. This configuration enables a fully injectable and minimally invasive deployment within deep tissues. Second, this work demonstrates for the first time that the PTE of DTCP can be increased by extending the implant length alone, without necessarily enlarging its cross-sectional area. Owing to its distributed coupling mechanism, DTCP also exhibits a high tolerance to lateral misalignment. These features together make DTCP particularly advantageous for human-scale spinal cord and PNS stimulation, allowing higher power delivery while maintaining minimal invasiveness. Third, the external DTCP driver is reduced to a compact, battery-powered signal generator and power amplifier, rendering the entire setup wearable and portable. Together, this combination of injectability, scalable efficiency, and portability establishes DTCP as a compelling foundation for the next generation of minimally invasive electroceutical systems. A summary comparison with representative state-of-the-art wireless neural stimulator systems is provided in Supplementary Table 1, outlining their operating frequency, powering modality, implant dimensions, and achieved power-transfer efficiency.

## Overview of DTCP-based injectable electroceuticals for nerve stimulation

Fig. 1b illustrates the conceptual design of the TINY system, shown here in the context of spinal nerve stimulation as an example application. Conventional spinal stimulation techniques typically rely on meter-long wired electrodes implanted subcutaneously through open-back surgery, with the effective stimulation region confined to the electrode tip. In contrast, the TINY measures around 1.5-2 cm in length and can be precisely injected into position using a customized needle during an outpatient procedure, significantly reducing invasiveness. The wearable battery-powered stimulator controller can be externally attached to the skin directly above the implant location.

The operating principle of DTCP is to establish an alternating E-field in tissue using two external transmitter (Tx) electrodes and to differentially harvest the resulting potential drop with two receiver (Rx) electrodes on the implant. When a sinusoid (typically tens of MHz) is applied across the Tx pair, displacement current traverses the high-impedance skin while conductive current dominates in the underlying muscle; together, these pathways form a closed loop through the body. The implant harvests the local field and converts the differential RF voltage into useful power.

To interpret coupling and guide design, the tissue volume between the two electrode pairs can be approximated by a layered lumped network (right panel in Fig. 1c). Each layer—skin, fat, and muscle—is represented by a complex vertical impedance (*Z*_y1_, *Z*_y2_, *Z*_y3_) and a transverse impedance (*Z*_x1_, *Z*_x2_, *Z*_x3_). The electrode–tissue interfaces are mainly capacitive at these frequencies, whereas bulk muscle exhibits higher conductance than skin or fat. Consequently, the delivered power at the implant depends on (i) the Tx spacing d_Tx_ relative to Rx spacing d_Rx_, (ii) implant depth and local layer thicknesses, and (iii) frequency, which jointly set the division of field lines. This model explains two key properties observed experimentally in later sections. (1) PTE increases with larger d_Rx_ even when d_Tx_ is increased proportionally. As d_Tx_ grows, the vertical tissue impedances (*Z*_y1_, *Z*_y2_, *Z*_y3_), set mainly by depth, change little, whereas the already large transverse impedances of skin and fat (*Z*_x1_, *Z*_x2_) grow further but contribute minimally to the net current path. The transverse muscle impedance *Z*_x3_, however, increases appreciably with d_Tx_, so a longer receiver pair samples a larger fraction of the field. (2) DTCP is tolerant to angular misalignment because the implant directly measures the potential difference at two points in tissue, avoiding dependence on coil orientation or an acoustic focal spot.

## Benchtop characterization of “TINY”

To evaluate the powering performance of the TINY implant, we establish a battery-isolated benchtop setup using differential RF excitation and real-time power monitoring (Fig. 2a). The entire system is powered by a floating battery supply to eliminate ground interference, and the single-ended RF signal from the generator is amplified and then split by a 180° –3 dB hybrid coupler into differential outputs. All subsequent connections use the same type and length of coaxial cables to ensure balanced transmission. Bidirectional couplers are inserted to measure forward and reflected power in real time, providing an accurate estimation of the overall system efficiency.

**Fig. 2.**
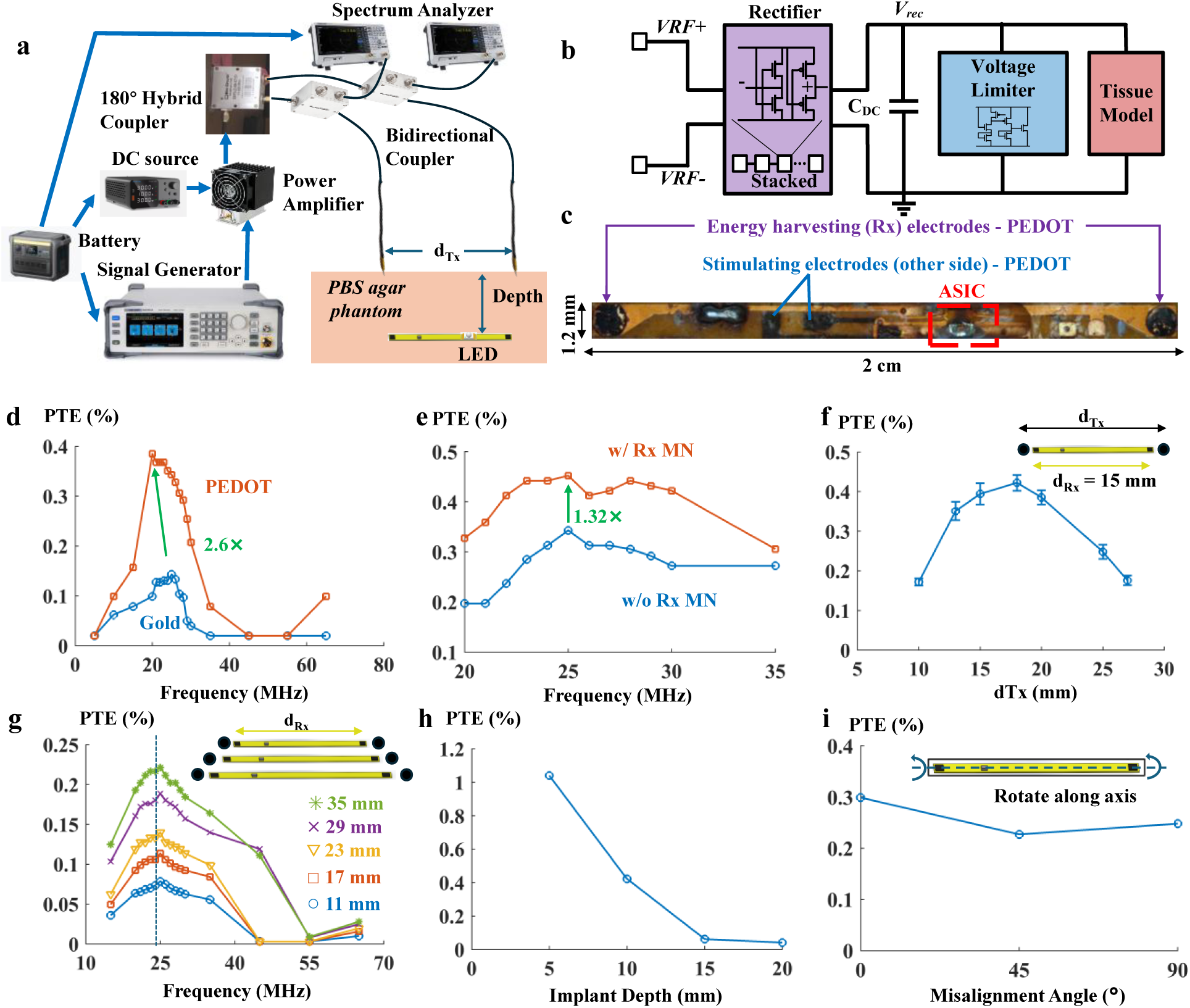
Benchtop characterization of TINY. **a.** Schematic of the bench test setup. A 180° –3 dB hybrid coupler converts the single-ended signal into a differential output, followed by equal-length coaxial cables for balanced transmission. Same bidirectional couplers monitor forward and reflected power in real time, enabling accurate estimation of the PTE. **b**. ASIC block diagram showing the impedance matching network, stacked cross-coupled rectifier, and voltage limiter for overvoltage protection. **c.** Photo of the implant with ASIC and PEDOT-coated energy-harvesting and stimulation electrodes. The device has a total volume of 2.7 mm3 and a weight of 9 mg. The LED is mounted on the stimulation electrodes for bench test only. **d**. PTE–frequency comparison of gold and PEDOT-coated surfaces. The PEDOT coating provides 2.6 × efficiency. **e.** Implant with matching network (MN) shows 32% higher efficiency than the one without. **f.** Best distance of Tx electrodes (dTx) for an implant with dRx=15 mm **g.** Plot showing that implants with longer distances between Rx electrodes (dRx) exhibit higher PTE. For each Rx length, the Tx location was optimized to achieve the best performance. **h**. PTE versus implant depth in phantom**. i.** PTE under rotation along the implant axis (0°–90°) shows only a maximum of 24% drop.

The implant’s wireless power reception is benchmarked using an integrated LED indicator. When the LED reaches its visible illumination threshold under dim indoor lighting, the net received power is measured to be 343 µW, which is used as the reference point for subsequent PTE evaluation. To assess performance under tissue-equivalent conditions, all bench measurements are conducted in a jelly-like phantom prepared by mixing 1× PBS diluted 20-fold with 1.8% agar. This composition yields an effective conductivity close to that of muscle tissue (~0.5 S/m) while providing stable and reproducible coupling for wireless power transfer characterization.

The custom-fabricated ASIC (Fig. 2b) integrates a rectifier and a voltage limiter. The cross-coupled differential drive (CCDD) rectifier is employed with a 4-stage cascaded structure to keep a balance between PTE and voltage conversion efficiency (VCE). A crowbar-style voltage limiter protects the circuits by rapidly sinking current once the supply exceeds a defined threshold. It will remain inactive under normal conditions to minimize power consumption. The detailed circuit implementation of the limiter is shown in Supplementary Fig. 1. As the stimulation mode represents the highest power demand, the rectifier is specifically sized and tuned for optimal efficiency under this operating condition. Its dimension after dicing is 0.6 × 0.3 × 0.3 mm.

Fig. 2c shows the implant consisting of the ASIC and PEDOT-coated energy-harvesting and stimulation electrodes. The device is 2 cm long and 1.2 mm wide on a 0.1 mm-thick flexible PCB, with a total volume of 2.7 mm^3^ and a mass of 9 mg. Because an optical indicator is unnecessary *in vivo* (light is not visible through tissue), the LED is omitted from the implant layout. Instead, for benchtop characterization, an LED is temporarily mounted across the stimulation electrodes to provide a visual readout of harvested power and stimulation activity.

To lower the Rx electrode impedance and improve the electrode–tissue contact, we electroplate PEDOT onto the gold electrodes. As shown in Fig. 2d, PEDOT coating increases PTE within the optimal frequency range by 2.6× relative to the uncoated electrodes, with noticeable gains across the entire measured band.

To further improve PTE, an impedance matching network (MN) is incorporated into the DTCP system for the first time. An L-type configuration with minimal components is implemented to balance performance and implant size. As shown in Fig. 2e, the MN improves the PTE by approximately 30%, compared to the unmatched case. Although theoretical modeling predicts much higher enhancement, the limited Q-factor (< 5) of the required high-value inductors (~5 µH at 25 MHz) introduced considerable loss, constraining the achievable gain. Moreover, the discrete components are larger in size than the ASIC, offsetting the benefits of miniaturization. These results indicate that, under the current design constraints, impedance matching provides measurable but modest improvement; however, its impact is expected to increase as system capacitance is further optimized in future miniaturized implementations.

We next analyze geometric parameters affecting PTE. For an implant with Rx electrode spacing (d_Rx_) of 15 mm at a depth of 10 mm (Fig. 2f), varying the transmitter spacing revealed an optimal d_Tx_ between 15 and 20 mm—slightly larger than d_Rx_. This can be explained by the nature of the dipole field distribution of differential electrodes: the E-field directly beneath each Tx electrode is relatively weak, while the strongest field region appears between the two Tx electrodes. When d_Tx_ becomes excessively large, tissue losses increase, and the Rx electrodes can no longer effectively harvest the full potential difference along the dominant field path. Therefore, the optimal d_Tx_ is typically slightly greater than d_Rx_.

Building upon the preceding geometric analysis, Fig. 2g compares the maximum achievable PTE across frequency for implants with different d_Rx_ at a depth of 10 mm. Importantly, d_Tx_ is not fixed but individually optimized for each d_Rx_ before comparison. Implants with a 35 mm length exhibit ~3× the PTE of the 11 mm implant, and this advantage persists across nearly the entire measured band. From the Rx electrodes to the ASIC, only a single wire needs to span the added distance, allowing the added width to be kept minimal. In our tests, the commercial flexible PCB is not precision-trimmed due to the vendor fabrication limits, but an in-house substrate [41] can readily achieve negligible area/volume growth while substantially extending length, thereby improving PTE at a minimum cost. This highlights a distinctive and previously unreported DTCP benefit: power delivery scales primarily with implant length, without a proportional increase in cross-section or overall volume.

Depth effects are also quantified. Fig. 2h shows the PTE of a 15 mm-long implant with PEDOT coating and MN optimization at different depths. The PTE reaches 1.03% at a depth of 5 mm and 0.42% at 10 mm. Beyond 15 mm, the efficiency degradation becomes more gradual.

Another major advantage of DTCP is its tolerance to misalignment. In Fig. 2i, rotating the Tx electrodes relative to the Rx from 0^∘^ to 90^∘^ reduces the minimum PTE by only 24%. This robustness arises because the DTCP Rx electrodes collect the potential difference directly between two contact points within the tissue. Therefore, as long as the field distribution at those two points remains unchanged, the implant’s efficiency is largely unaffected by its orientation. In contrast, for coil-based links, the efficiency at 90° typically approaches zero, and ultrasonic links also suffer from narrow alignment windows. This comparison highlights the superior misalignment tolerance of DTCP technology.

## Portable transmitter system

To enable a fully wearable system suitable for outpatient neural stimulation, we developed a battery-powered wireless transmitter prototype. Fig. 3a outlines the architecture: an MCU (MSPM0C1104) programs a direct digital synthesis (DDS) chip (AD9850) to generate a low-power RF tone with MCU-controlled frequency and amplitude (the latter also adjustable via an onboard potentiometer). The battery output is boosted by an LTC3122 converter to supply an RF power amplifier (PMA3-43-1W) that provides approximately 22 dB of gain. The amplified signal then passes through a balun (MABAES0060) to generate a differential output driving two Tx electrodes. As shown in Fig. 3b, the fabricated prototype measures 63 mm × 38 mm and is powered by a flexible 3.7 V lithium battery. Applying a pulsed drive to the power amplifier (500 µs/s) yields an average system power consumption of 0.29 W, while both pulse width and stimulation frequency are fully programmable via the MCU for flexible protocol control. The system attains a maximum output power of 28 dBm (Fig. 3c; measured with a 20 dB attenuator on the spectrum analyzer). Although fabricated on FR-4, the same design can be migrated to a flexible PCB to enhance wearability.

**Fig.3.**
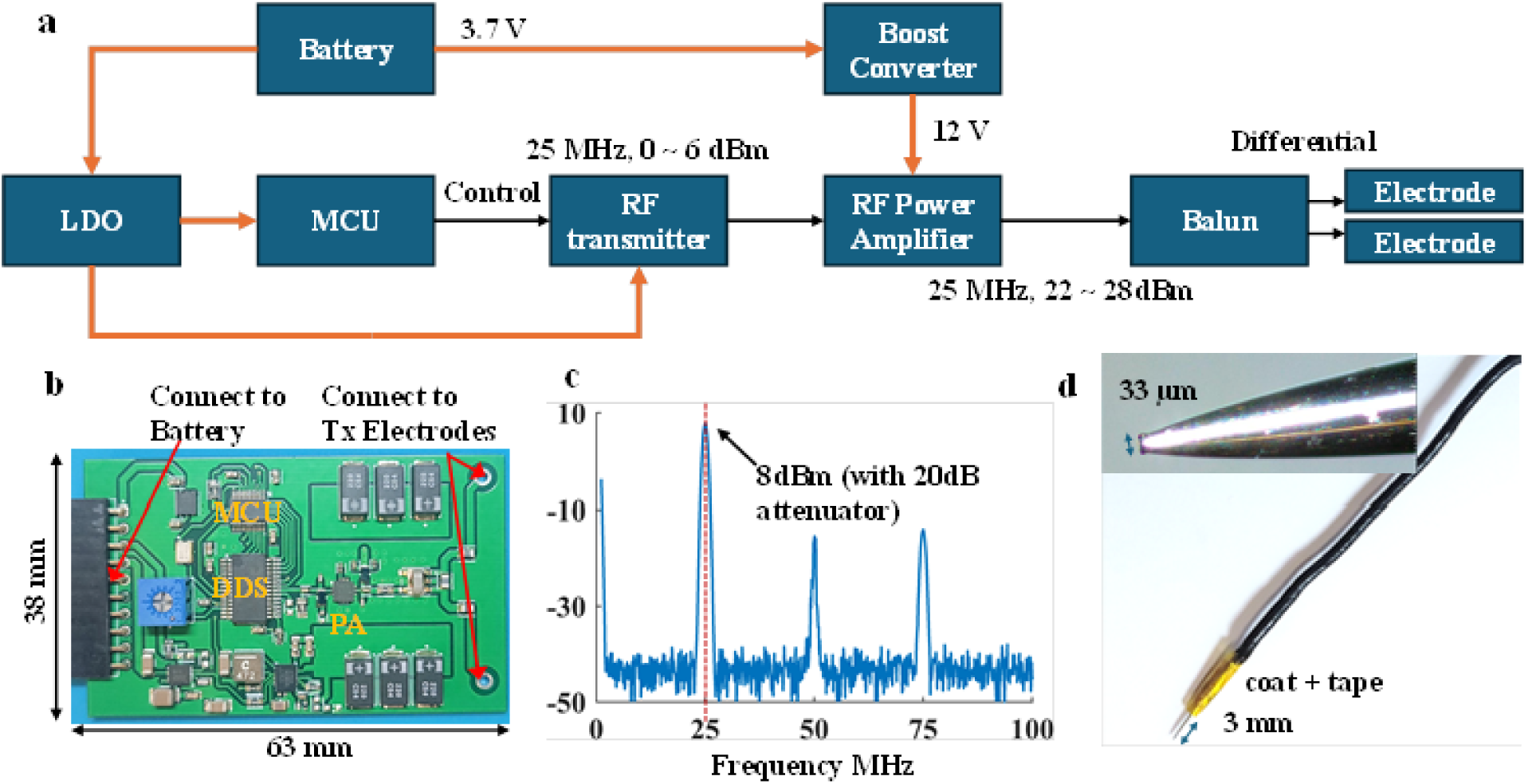
Portable DTCP transmitter system for wireless powering of injectable implants. **a.** System block diagram of the battery-powered wireless transmitter. The circuit integrates an MCU-controlled DDS to generate a programmable RF carrier, followed by a boost converter and RF power amplifier to drive a balun-connected differential output for the external Tx electrodes. **b.** Photograph of the fabricated prototype (63 mm × 38 mm) that operates from a 3.7 V lithium battery and supports programmable pulse-modulated operation. **c.** Measured RF output spectrum demonstrating a maximum transmitted power of 28 dBm at fundamental frequency (with a 20 dB attenuator). **d.** Micro-pin Tx electrode designed to enhance tissue coupling with minimal invasiveness. The stainless-steel electrode features a 33 µm tip width, 3 mm exposed length, and 330 µm maximum diameter at the base end—smaller than typical CGM needles—coated with PDMS and wrapped with polyimide for insulation and robustness.

The Tx electrodes, shown in Fig. 3d, are designed to improve PTE while minimizing invasiveness. Because the conductivity of skin is low [39], significant transmission loss occurs through these layers [30]. To address this, a micro-pin stainless-steel electrode is developed to penetrate the skin and enhance coupling efficiency with minimal tissue disruption. The pin features a 33 µm tip width, 3 mm exposed length, and 330 µm maximum diameter, smaller than commercial continuous glucose-monitoring (CGM) needles, indicating that, while minimally invasive, its tissue impact is limited. The proximal end of the pin is press-fit into a copper lead and soldered at the junction. The remaining exposed portion is coated with PDMS and wrapped with polyimide tape to provide waterproofing and maintain mechanical robustness during implantation. Additional benchtop measurements using non-invasive surface electrodes are presented in Supplementary Fig. 2. While surface coupling is feasible, stable fixation on rat skin proves challenging and limits repeatability; therefore, pin electrodes are adopted in the main experiments to ensure repeatable conditions.

According to the IEEE C.95.1 safety standard, the maximum permissible current for human contact at 25 MHz is 31.5 mA [38]. The electrode–tissue impedance measured on a rat hindlimb using a VNA ranges from 325 Ω to 362 Ω in resistance and from 27.2 pF to 38.6 pF in capacitance. These values vary due to differences in electrode–tissue contact and micromotion from respiration. With 28 dBm of RF power, considering a stimulation duty cycle of 500 µs/s, the resulting RMS current is only 0.61 mA, which is well below the safety limit specified by the standard. A detailed derivation of the corresponding current and power levels is provided in Supplementary Note 1.

## TINY system demonstrates wireless peripheral nerve stimulation

To demonstrate peripheral nerve stimulation and assess the potential for clinical translation, the TINY system is implanted in the rat hindlimb and wirelessly driven by the portable transmitter. The wirelessly powered implant reliably evokes compound muscle action potentials (CMAPs) and visible leg kicks when placed near the sciatic nerve, successfully achieving direct in vivo stimulation of peripheral nerves (n = 3).

Stimulation in a rat with the TINY system, wirelessly powered at an 8 mm Tx–Rx separation, is shown in Fig. 4a. The implant was positioned between muscle and nerve with the stimulation electrodes facing the nerve. A 500-µs monophasic pulse train at 1 Hz was applied from the portable transmitter (25 MHz, 28 dBm RF power). Average EMG recordings from the biceps femoris muscle show responses time-locked to the applied stimuli (t=0 s in Fig.4a) at the same frequency. To exclude direct stimulation by the external field, we performed a negative control using the transmitter alone at the same power; only stimulus-locked artifacts were observed, with no evoked EMG responses. The stimulation capability and the dependence on electrode geometry are further analyzed through finite-element modelling, as detailed in Supplementary Fig. 3

**Fig.4.**
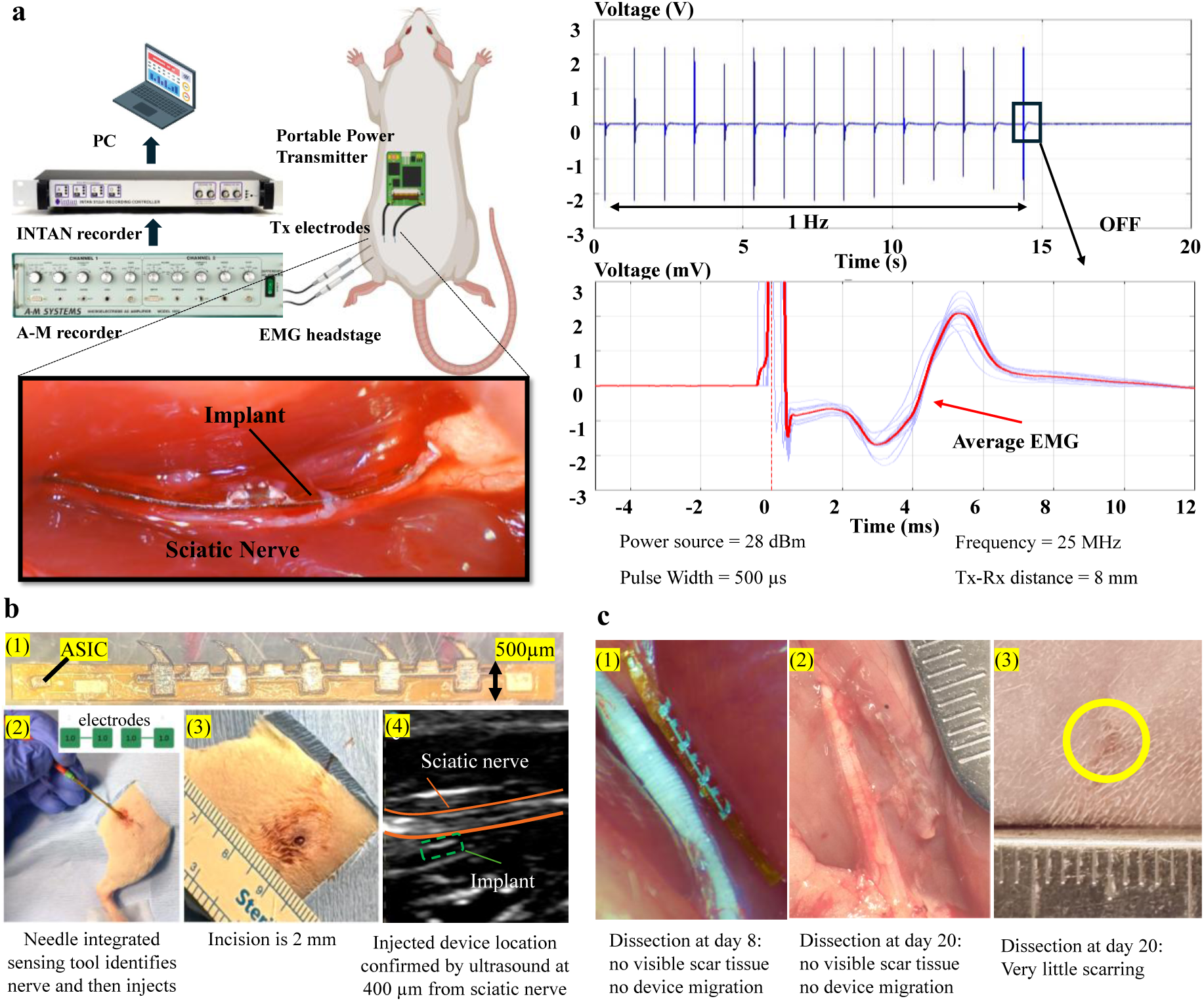
In vivo demonstration of wireless peripheral nerve stimulation and post-injection stability of the TINY system. **a.** Experimental setup for wireless sciatic nerve stimulation in rats using the portable DTCP transmitter. The transmitter drives two external Tx electrodes while EMG signals are recorded from the biceps femoris muscle. The implant is positioned adjacent to the sciatic nerve. Wireless stimulation evokes repeatable CMAPs at 1 Hz. Representative EMG trace (top) and averaged response (bottom) show time-locked muscle activation synchronized with each stimulus pulse (500 µs width). **b.** Injection of the miniaturized TINY dummy prototypes for migration studies. (1) Optical micrograph of the 500 µm device with biological anchors to enhance long-term positional stability (2) Integration of the nerve sensor into a 20-gauge spinal needle for injection. (3) Photograph of a 2 mm skin incision prior to injection. (4) Ultrasound imaging confirming device placement (green) adjacent to the sciatic nerve (orange). **c.** Post-implantation tissue images demonstrating long-term positional stability and minimal tissue response. (1) 8 days after injection, the implant remains at the intended location without migration or encapsulation. (2) At 20 days, the implant continues to reside near the nerve with no visible fibrotic reaction. (3) External skin inspection at day 20 shows only a small residual mark (yellow circle), indicating minimal invasiveness.

## Migration and damage study of TINY after injection

The purpose of this section is to investigate the longitudinal implant migration in rats to assess positional stability of TINY after injection. We fabricate 500 μm-wide flexible wireless neurostimulator prototypes incorporating ASICs, though their functionality is not evaluated in this study (Fig. 4b-1). The 1.2 mm-wide prototypes used for benchtop characterization reflect current PCB fabrication limits; with optimized in-house processes, the design is expected to scale down to 500 μm without altering functionality. Surface features are engineered to promote localized scarring, serving as biological anchors to enhance long-term positional stability. In our rat study (n=4), we inject the devices parallel to the branches of the sciatic nerve using a nerve-sensing probe integrated into a 20-gauge spinal needle via a 2 mm incision (Fig. 4b-2,3). After positioning the spinal needle near the nerve, it advances deeper into the tissue and aligns parallel to the nerve with the assistance of the nerve-sensing probe. In Fig. 4b-2, the four green pads on the probe are two pair of sensing electrodes used to detect nerve proximity; the numbers “1.0” within the pads indicate the model-predicted probability of nerve tissue with data processing supported by deep learning. Saline hydro dissection is performed to create a pocket adjacent to the nerve for the chip. The chip is then injected into the pocket parallel to the nerve using either (a) a push rod or (b) alginate hydrogel. After the chip is implanted, the needle is carefully withdrawn while injecting 1 mL of hydrogel along the needle trajectory. Ultrasound imaging confirms the successful implantation of the chip adjacent to the sciatic nerve (Fig. 4b-4).

Tissue dissection conducted at 8- and 20-days post-implantation confirms the precise localization of the implanted device, with no observable signs of migration or fibrotic tissue response, as shown in Fig. 4c. On day 8 (Fig. 4c-1), the device remains securely positioned at the targeted implantation site, with no visible displacement or tissue encapsulation. By day 20 (Fig. 4c-2), the implant continues to maintain its position adjacent to the nerve, further supporting the long-term mechanical stability of the device. Notably, no macroscopic scar tissue is detected at either point, indicating minimal chronic inflammatory response. Additionally, examination of the external skin at the injection site on day 20 (Fig. 4c-3) reveals only minimal surface scarring, indicating that the implantation method is both minimally invasive and well-tolerated. These results collectively validate the stable in vivo positioning of the device and its biocompatibility over a 20-day period. All surgical procedures are approved by the Institutional Animal Care and Use Committees (IACUC) of the University of Florida and Harvard University.

## Discussion

This work presents the first fully injectable neural stimulator “TINY” based on DTCP, which is wirelessly driven by a compact, wearable transmitter. TINY integrates the receiver electrodes, ASIC, and stimulation electrodes into a flexible strand deliverable through a fine needle, enabling minimally invasive electroceuticals. Leveraging the unique coupling mechanism of DTCP, TINY can maintain a scalable PTE that increases with implant length without requiring a larger cross-section, substantially reducing the cost of achieving higher PTE in applications demanding greater power. TINY also exhibits strong tolerance to angular misalignment between the external and implanted electrodes. Considering that the implant may shift or rotate within tissue, users can simply reposition the external electrodes to find the optimal coupling, whereas other WPT methods might require surgical realignment. Combined with the battery-powered portable transmitter, these features enable practical and flexible use of TINY for future distributed and wearable neurostimulation applications. This technological architecture readily supports clinical translation and may ultimately permit patients to operate implants safely in home or outpatient settings.

For a proof-of-concept demonstration, TINY is implanted in the rat hindlimb, where direct electrical stimulation reliably evokes both neural responses (EMG) and visible leg movements. The portable, battery-powered transmitter delivers approximately 630 mW of RF power in vivo to operate TINY, and the resulting RMS contact current is 0.61 mA, which remains well below the IEEE [38] and ICNIRP [42] safety limits. Even during transmission bursts, the instantaneous peak current stays within the permissible range, indicating a wide safety margin that supports future integration of additional functionalities such as neural recording or biochemical sensing. The data we provide serve as evidence that TINY can stimulate the sciatic nerve through wireless power. Additional work is still needed to advance this technology toward a biomedical device suitable for clinical use.

First, to achieve high transmission efficiency in the current setup, we employ a micro-pin type Tx electrode similar in concept to those used in CGMs. Although the dimension is smaller, this configuration remains invasive. At the human scale, however, optimized Rx electrode designs— such as 3D structures or improved surface coatings to enhance tissue contact—together with better impedance matching on both Tx and Rx sides can collectively raise the overall PTE. With sufficient PTE, effective neural stimulation can be achieved using surface-mounted external Tx electrodes, allowing the transmitter system to operate in a fully non-invasive manner.

Second, the present prototype still has significant potential for further miniaturization, as a large portion of the substrate does not contribute to PTE due to the precision constraints of commercial PCB fabrication. With in-house micro-fabrication enabling thinner and narrower substrates, and an ASIC designed with adaptive impedance matching, TINY will achieve additional miniaturization, further reducing the surgical burden.

Third, the long-term biocompatibility of TINY requires further evaluation. The current prototype uses copper traces coated with polyimide and gold, and potential copper-ion leakage over prolonged implantation remains to be investigated. Similar to the previous point, in-house fabrication using Pt or other inert materials for trace formation can eliminate this concern.

Finally, the origin of the optimal operating frequency of TINY remains unclear, as it results from a complex coupling among circuit behavior, electrochemical interface, and electromagnetic field distribution. Understanding this mechanism will be valuable for refining the matching network design and further improving overall system efficiency.

In summary, the DTCP technology and the TINY system introduced in this work open multiple opportunities for future electroceutical research and clinical translation. The core concept of DTCP, using tissue as the conductive and capacitive medium for power delivery, eliminates the need for tightly coupled coils or acoustic channels that often limit alignment tolerance and miniaturization.

This principle provides a broadly applicable powering and communication approach for diverse neural interfaces across the nervous system. Beyond sciatic nerve stimulation shown in this paper, the same principle can be applied to spinal cord and dorsal root stimulation for pain and motor rehabilitation, as well as to peripheral targets such as vagus, sacral, and tibial nerves for regulating inflammatory, gastrointestinal, and urinary functions. Moreover, TINY could be adapted for somatic nerve interfaces that restore motor and sensory functions in paralyzed or injured limbs, or for distributed sensor–stimulator networks enabling closed-loop neuromodulation. By supporting fully injectable, flexible, and wearable-compatible architectures, TINY establishes a new foundation for minimally invasive electroceutical systems that bridge experimental neuroengineering and future clinical medicine.

## Methods

### Wireless PTE characterization

To quantify the energy harvested by the implant during the experiment, a conventional approach would be to connect it through lead wires. However, in the case of DTCP, any such wires would extend well beyond the implant and could significantly disturb the surrounding E-field, introduce additional variables, and compromise measurement accuracy. Therefore, an LED load is integrated onto the implant to enable wireless PTE measurement in bench tests, a feature particularly beneficial at MHz frequencies. First, the implant is characterized using a vector network analyzer (VNA) to measure the input-side S_11_. Then, the input terminal is connected to a signal generator, and the input power gradually increases until the LED on the implant reaches its visible illumination threshold under dim indoor lighting. The corresponding input power at this boundary condition is recorded. By subtracting the reflected power from the total input, the net power delivered into the implant is calculated to be 343 µW. Therefore, in the subsequent bench tests, whenever the LED again reaches the same illumination threshold, it is assumed that the implant received 343 µW of power, which is then used to calculate the overall PTE of the system.

The phantom used in the experiments is prepared by mixing 1× PBS diluted 20-fold with 1.8% agar, forming a jelly-like tissue-mimicking medium. This specific formulation is chosen for several reasons. First, using undiluted PBS would lead to unrealistic conditions, since the conductivity of muscle tissue is around 0.5 S/m [39], whereas that of standard PBS is about 11 S/m [40], far higher than physiological levels. Second, employing a purely liquid medium would result in perfect and constant contact between the phantom and the implant, leading to an overly optimistic estimation of PTE. To address these issues, PBS is diluted to reduce conductivity, and agar is added to solidify the mixture. Some studies use biological tissues such as pork or ground beef as phantoms; however, once exposed to air, their surface moisture changes within minutes, introducing uncontrolled variability. Moreover, since an LED-based optical indication is used to assess PTE, the color and opacity of real tissue could interfere with optical detection.

### Impedance matching network design

The impedance MN was designed based on in vivo impedance measurements of the electrode–tissue interface. A dummy implant with the same PEDOT-coated electrodes but without internal circuitry was used to measure the electrode impedance in vivo using a vector network analyzer (VNA). Subsequently, the input impedance from the electrodes to the ASIC terminals was characterized under identical settings. Using these two measured impedance values, an L-type MN topology was modeled and optimized in simulation to maximize PTE. This configuration was selected because it has only two passive components, offering the simplest structure and smallest footprint. In practice, the required inductance for impedance matching was on the order of 5 µH due to the small system capacitance and the resulting high impedance level. At 25 MHz, commercial 0603 or 0402 inductors of this value exhibited a low Q-factor (< 5), introducing substantial loss. Reducing the inductance to achieve a higher Q-factor shifted the matching frequency and diminished the matching effect. After several combinations and iterations, the best experimental configuration yielded a ~30% increase in PTE as shown in Fig. 2e. However, the added discrete components increased the total volume from 2.7 mm^3^ to 3.4 mm^3^, thus limiting the benefit for miniaturized systems. These trade-offs will be revisited as future designs optimize the system capacitance and integration level.

### TINY implant fabrication and assembly

The ASIC was fabricated using the TSMC 65 nm RF LP process with a 1P9M + Al metal stack, consisting of nine copper layers and a top aluminum layer serving as the exposed pad metal. After wafer processing, the die was diced using a 15 µm-wide diamond blade, resulting in individual chips measuring 0.6 × 0.3 × 0.3 mm^3^ (length × width × thickness). The die can be further thinned to reduce thickness for future implant versions.

The flexible substrate was a 100 µm-thick polyimide PCB with electroless nickel immersion gold (ENIG) surface finish, featuring a 3 µm gold plating. The ASIC was picked and placed using a T-4909-AE manual die bonder and attached onto the PCB with MG Chemicals 9310 non-conductive epoxy, cured at 120 °C for 10 min. Prior to wire bonding, both the die surface and PCB pads were cleaned sequentially with acetone and deionized water. K&S 4124 Ball Bonder was employed for gold wire bonding. Because the flexible substrate provides limited mechanical rigidity, the wedge-bond joints were reinforced using EPO-TEK® H20E silver epoxy, cured at 120 °C for 15 min. The entire assembly was then encapsulated with SYLGARD™ 184 PDMS (10:1 base to curing agent). After degassing for 10 min, the PDMS was applied uniformly over the ASIC and bonding wires using a micro-pin tool, followed by curing at 150 °C for 10 min. This coating procedure was repeated multiple times until sufficient encapsulation thickness was achieved to fully cover the bonded area.

The implant’s Rx electrodes measured 900 × 900 µm^2^, while the stimulation electrodes were 600 × 600 µm^2^ with a 1.4 mm pitch. To reduce electrode impedance and enhance tissue contact, PEDOT electroplating was performed. The plating solution consisted of lithium perchlorate (LiClO_4_, 120 mM, 0.24 g) and EDOT monomer (30 mM, 60 µL) dissolved in 20 mL acetonitrile (ACN). A stainless-steel wire (diameter 254 µm) served as the negative electrode, and deposition was conducted using a DC current of 100 µA for 100 s. After coating, the electrodes were rinsed sequentially with acetone and deionized water. For devices incorporating an impedance matching network, 183°C solder was used for component attachment, and the soldered area was subsequently covered with the same PDMS formulation for protection and insulation.

### Phantom preparation and benchtop characterization

The phantom was prepared by mixing 1× PBS diluted 20-fold with 1.8% agar powder (w/v). The mixture was stirred and microwaved alternately until the solution became clear and homogeneous, ensuring complete dissolution of agar. The liquid was then refrigerated at 4 °C for 2 h to allow solidification, resulting in a transparent and uniform gel phantom suitable for electrical characterization.

All benchtop measurements were conducted using a fully battery-powered setup to eliminate ground loops and minimize electromagnetic interference. A 25 MHz sinusoidal signal was generated (Siglent SSG3021X), amplified ( Mini-Circuits ZHL-20W-13+) and then fed into a 180° –3 dB hybrid coupler (Mini-Circuits ZAPDJ2-5W-521S+) to produce a balanced differential signal.

Each output arm of the coupler was connected through identical SMA coaxial cables to a bidirectional coupler (Mini-Circuits ZFBDC26-52HP-S+), enabling simultaneous monitoring of forward and reflected power. During testing, Siglent SSA3032X spectrum analyzer was used to measure both forward and reverse power from the couplers. The net power delivered into the phantom was calculated as the sum of forward–minus–reflected power from both channels and then used to determine the system PTE.

### Surgery Technique

Adult male Sprague–Dawley rats (400–500 g) were anesthetized with 2–3% isoflurane in oxygen (500 ml min^⁻¹^) and maintained at 36–37 °C using a thermostatically controlled heating pad. The tail was grounded, and the left thigh was shaved and sterilized. A lateral skin incision was made along the left hind limb, extending from the hip to the knee, parallel to the femur. The vastus lateralis and biceps femoris muscles were gently separated to expose the sciatic nerve. A wireless stimulator was positioned with its stimulation electrodes aligned to the nerve and its power electrodes facing the vastus lateralis muscle. The skin incision was then closed with 4-0 polypropylene sutures. The transmitter circuit and electrode fabrication were described in detail in the main text (Fig. 3). After closure, two transmitter electrodes (micro-pins) were inserted percutaneously, one near the hip and the other near the knee, approximately 22-25 mm apart. At the end of the experiment, animals were euthanized according to approved protocols. All procedures were approved by the Institutional Animal Care and Use Committee of the University of Florida (IACUC) and conducted in compliance with institutional and NIH guidelines.

### Data acquisition and analysis

EMG recordings were obtained using TE/S46-638 Pt/Ir subdermal needle electrodes. The signal and reference electrodes were inserted into the biceps femoris with an inter-electrode spacing of 8 mm, and the ground electrode was placed on the rat’s neck. Signals were amplified by an AM1800 recorder (bandwidth 0.5–5 kHz, gain 1000×) and digitized by an Intan recording system at a 30 kHz sampling rate. Data were extracted using MATLAB scripts. Stimulation was applied with a pulse pattern of 500 µs width at 1 Hz. Each stimulation artifact was used to synchronize the EMG responses across trials. The signals were segmented and averaged across 30 repetitions to obtain the mean EMG waveform shown in Fig. 4a.

## Data availability

The dataset that supports the plots within this paper and other findings of this study are available in the Supplementary Information and from the corresponding author upon reasonable request.

## Code availability

Custom-developed MATLAB codes for extraction and plotting are available from the corresponding author upon reasonable request.

